# Splicing-guaranteed artificial introns enable efficient homology-directed repair at internal exons and streamlined cell engineering

**DOI:** 10.64898/2026.10.16.759525

**Authors:** Shang Wang, Xianteng Wang, Mengyu Shang, Kai Wu, Mengkai Zhang, Weiren Huang

## Abstract

Selection-marker-assisted homology-directed repair (HDR) efficiently edits coding sequence termini but is poorly suited for internal exons. Here, we developed artificial reporter intron-based HDR (ARI-HDR), which inserts a selectable intron into target exons to enable precise internal-exon editing. Although designed for scarless removal by splicing, ARIs frequently exhibit inaccurate splicing. Using a random mutagenesis reporter assay, we identified intron upstream sequences (IUS) as key determinants of splicing fidelity and screened degenerate IUS variants to obtain reliably spliced ARI donors. ARI-HDR successfully introduced site-specific mutations into five human genes. Combining long single-stranded DNA donors with conversion-tract optimization increased editing efficiency to over 80%. This efficiency enabled “Dirty-Editing,” a streamlined workflow that generates precisely edited mixed-cell populations without single-cell cloning. Together, ARI-HDR extends selection-based enrichment to internal exons across protein-coding genes, while our splicing-optimized intron design offers a broadly applicable solution for artificial intron-based technologies.

## Background

The promise of homology-directed repair (HDR) for precise genome editing is fundamentally constrained by its low efficiency (*1–3*). Current enrichment strategies rely on selection-marker labeling, but these are largely confined to the termini of coding sequences (CDS), leaving the vast majority of disease-relevant mutations in internal exons inaccessible (*4–6*). Even when editing succeeds, the residual exogenous sequences in the final transcript risk perturbing gene regulation (*7*).

Introns offer an elegant solution: they can carry functional cargo yet are excised from the mRNA (*8*). Artificial introns, designed for precise self-excision, have been proposed as a universal vehicle (*9*), but their broader application has been thwarted by the inherent complexity of RNA splicing. This field lacks a mechanistic understanding of why artificial introns frequently fail to splice correctly, and consequently, a reliable design strategy remains elusive (*10*).

Here, we present a systematic solution to this problem. We developed an artificial reporter intron (ARI) that functions as a portable, splicing-guaranteed selection marker, capable of enriching edits at any internal exon. By dissecting the molecular causes of aberrant splicing—namely, the influence of intron upstream sequences and cryptic splice site activation, we established design principles that ensure splicing accuracy. Coupled with optimized single-stranded DNA donors and minimized conversion tracts, this ARI-HDR platform achieves editing efficiencies exceeding 80% in human cells. This level of performance enabled us to bypass single-cell cloning entirely, establishing a streamlined “Dirty-Editing” workflow that rapidly yields mixed populations of precisely edited cells. Our work provides a comprehensive, efficient, and reliable solution for HDR-based editing, overcoming limitations in locus targeting, splicing fidelity, and experimental throughput. Unlike SMArT, which uses a transient AND-gate reporter on the HDR donor to enrich correct integrations (*11*), ARI-HDR attacks a complementary problem: extending selection-marker labeling to internal exons while leaving no exogenous sequence in the mature transcript.

## Results

### Overview of ARI-HDR gene editing strategy

Introns are non-coding genomic sequences that are excised from pre-mRNA by the spliceosome during splicing. They harbor regulatory elements that enable alternative splicing and modulate gene expression (*12, 13*). To develop an artificial intron-based editing strategy, we synthesized a universal artificial reporter intron (ARI) flanked by splicing donor and acceptor sites to ensure proper splicing, and carrying a selectable marker. The ARI serves as an HDR donor that splits exons and labels edited alleles; its excision during splicing ensures a scarless mature mRNA.

We proposed the ARI-HDR strategy as follows:

1. Cas9/gRNA generates a double-strand break (DSB) within an exon.
2. Homology arms (HAs) containing the ARI serve as donors for HDR-mediated DSB repair.
3. Desired mutations are introduced via the homology arms.

The edited gene thus contains both the ARI and the desired mutation. The selectable marker within the ARI enables enrichment of edited cells. Upon transcription, the ARI is excised along with native introns during pre-mRNA splicing, leaving a scarless mature mRNA **(Fig. 1A)**.

**Figure 1.**
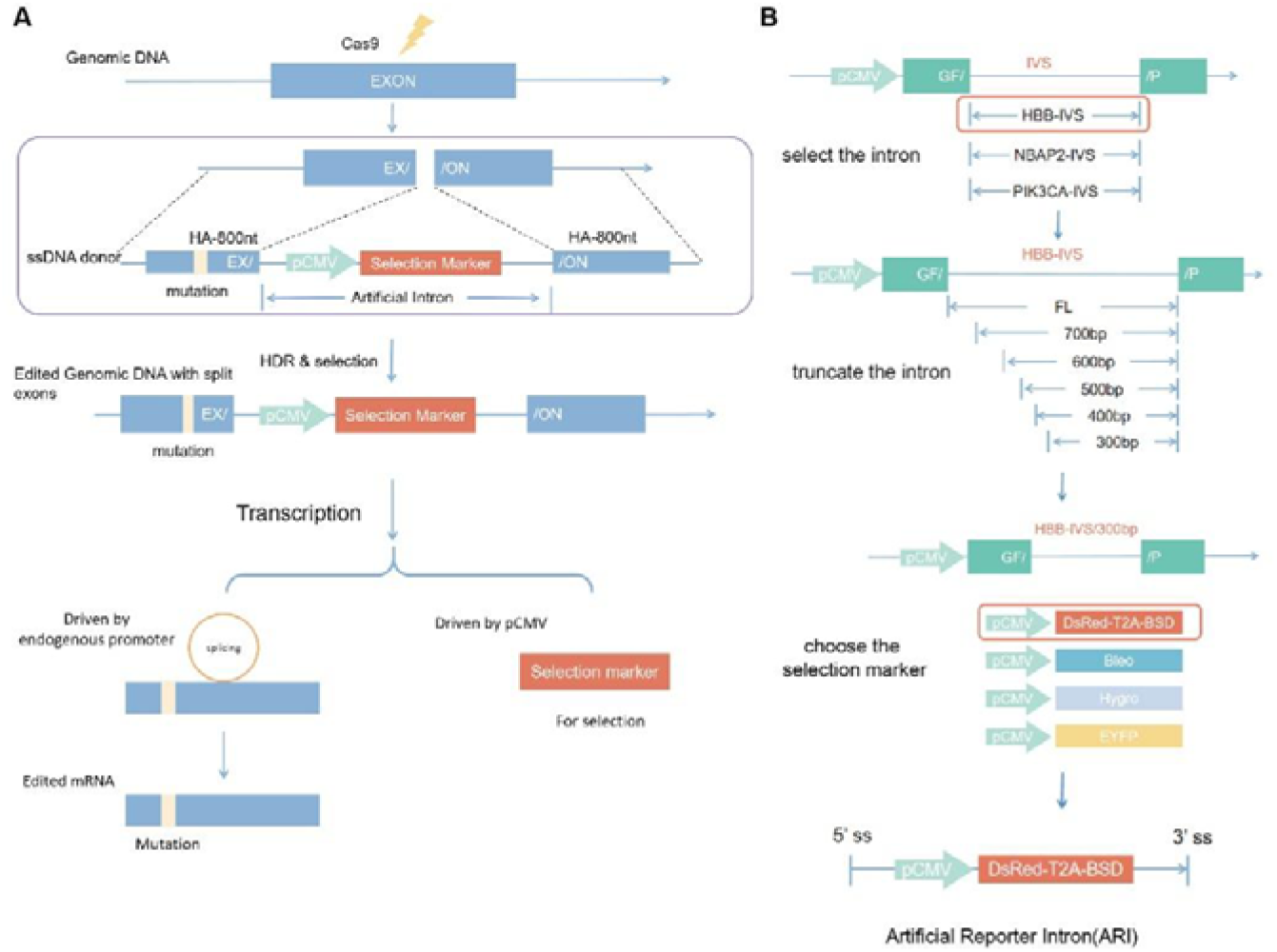
A. Overview of the ARI-HDR gene editing strategy. A DNA donor template carrying both the desired point mutation and an artificial reporter intron (ARI) was co-transfected with CRISPR/Cas9 components into target cells. Upon successful homology-directed repair (HDR), the donor template is integrated into the genome, generating an edited allele that contains both the mutation and the ARI. The ARI expresses a fluorescent marker enabling fluorescence-activated cell sorting (FACS)-based enrichment. The endogenous splicing machinery excises the intronic sequences from the target gene transcript, producing mature mutant mRNA that retains normal coding potential and ensures proper protein function. B. Optimization of the artificial reporter intron using a GFP-expressing vector. Three sequential optimization steps were performed: first, screening of three candidate natural introns derived from human *HBB*, *NABP2*, and *PIK3CA*genes; second, progressive 5′ truncation of the selected intron in 100 bp increments; third, insertion of various reporter genes into the artificial intron to identify the optimal configuration.

### Optimization of Selection Marker-Embedded Artificial Reporter Intron (ARI)

To develop the desired ARI, we first constructed a CMV-driven CopGFP reporter plasmid. Candidate introns derived from human *HBB*, *PIK3CA*, *NABP2* genes were inserted into the GFP CDS for mRNA and protein expression in HEK293T cells. All three introns mediated correct splicing and produced full-length GFP protein, with the *HBB*-derived intron (HBB-IVS) yielding the highest mRNA expression (**Fig. S1 A-D**). We then progressively truncated the HBB-IVS from the 5′ splice donor site in 100-bp increments. An intron as short as 300 bp (HBB-IVS/300) retained correct splicing activity (**Fig. S1 F-G**). To identify an optimal reporter gene for the Artificial Reporter Intron (ARI), we inserted different markers the HBB/IVS-300 intron. RT-PCR analysis revealed that only DsRed-T2A-BSD maintained correct splicing (**Fig. S1 H-K**).

Given its splicing fidelity (no aberrant variants) and robust reporter output, we selected the DsRed-T2A-BSD-embedded HBB/IVS-300 as the final ARI for subsequent studies (**Fig. 1B**).

### G-ARI-N rule is required for correct splicing

To determine the requirements for accurate ARI splicing, we analyzed the sequence preferences of human and mouse introns. Both human and mouse introns exhibited a strong preference for guanine (G) at the −1 position of the 5′ splice site (5′ss), whereas the 3′ splice site (3′ss) showed no clear nucleotide preference (**Fig. 2A, S2A**).

**Figure 2.**
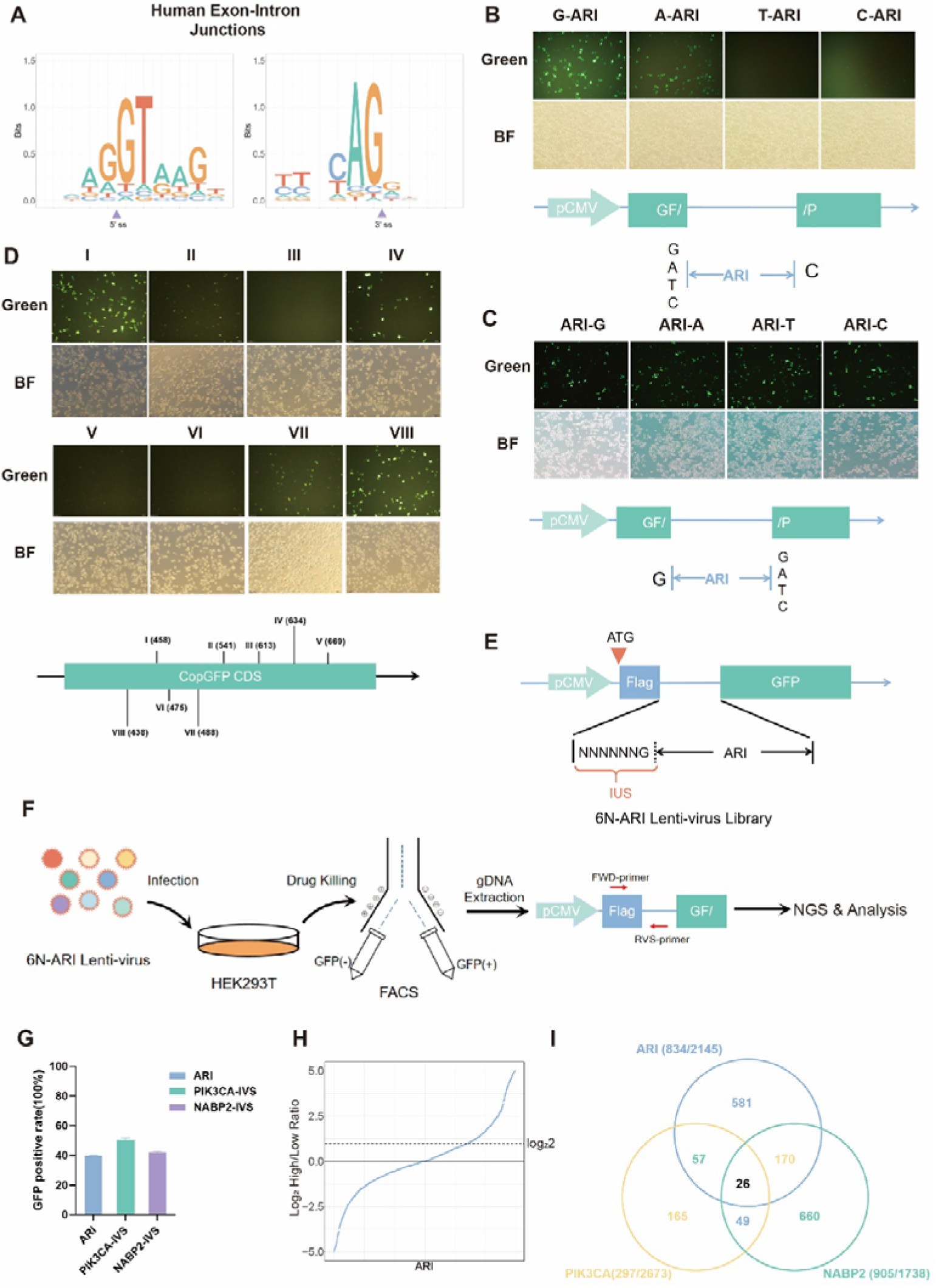
A. All human intron/exon junction sequences were analyzed. The 5’ splice site (left) shows strong nucleotide conservation with a canonical "GU" dinucleotide at the intron start, preceded by enriched "G" in the upstream exon sequence. In contrast, the 3’ splice site (right) is marked by the conserved "AG" motif at the intron end, with no typical sequence preference found in the downstream exon region beyond this boundary (as indicated by low information content). Y-axis represents information content (bits); purple arrowheads denote the exact exon-intron boundaries. B, C. Expression of a CMV-driven GFP open reading frame (ORF) disrupted by ARI insertion, with degenerate mutations introduced in the 5′ (B) and 3′ (C) flanking regions. Fluorescence microscopy (Green) and bright-field (BF) images show GFP expression levels across varying flanking sequences (B: G/A/T/C-ARI; C: ARI-G/A/T/C). Corresponding schematic diagrams illustrate the pCMV-GFP construct with ARI disrupting the ORF; functional splicing restores the GFP reading frame, with fluorescence intensity dependent on the specific 5′ or 3′ flanking bases. D. Insertion of ARI at eight different loci following the G-ARI-N rule disrupts GFP expression. Fluorescence (Green) and bright-field (BF) microscopy show varied GFP restoration across loci I–VIII within the CopGFP CDS. Loci I, IV, VII, and VIII exhibit robust GFP fluorescence, whereas II, III, V, and VI show minimal to no expression, indicating position-dependent splicing efficiency or ORF disruption despite the conserved flanking rule. E. Schematic of the reporter construct. A Flag-GFP fusion coding sequence (CDS) was interrupted by a random hexanucleotide sequence (NNNNNN) followed by G-ARI. Correct splicing of ARI, facilitated by specific 6N sequences (labeled as iUS), is required for in-frame translation of functional GFP. The construct is driven by the pCMV promoter with an ATG start codon upstream of the Flag tag, and is utilized for the 6N-ARI Lenti-virus Library. F. Workflow for Screening 6N Sequences that promote correct ARI splicing. HEK293T cells were infected with the 6N-ARI Lenti-virus library, followed by drug killing and FACS sorting into GFP (-) and GFP (+) populations. Genomic DNA (gDNA) was extracted and the target region (pCMV-Flag-ARI-GFP) was amplified using FWD/RVS primers, followed by NGS & analysis. G. Flow cytometry results of GFP positive cells of puromycin killed library lenti-virus infected HEK293T cells. The bar chart quantifies the GFP positive rate (%), with data presented as mean ± SD from three independent experiments (n=3). H. Rank plot of log□-transformed enrichment scores for each hexanucleotide (NNNNNN) motif derived from the screening data (F). The enrichment score is calculated as log□ (counts in GFP cells / counts in GFP cells). Motifs are ranked in ascending order of enrichment score on the x axis; the y axis indicates log□ fold enrichment. Higher values denote motifs preferentially associated with productive ARI splicing. I. Venn diagram illustrating the overlap among 6N sequence motifs that promote splicing of three different introns. The diagram quantifies motif distributions for ARI (834/2145), PIK3CA (297/2673), and NABP2 (905/1738), with unique counts of 581, 165, and 660, respectively. Pairwise overlaps include 57 (ARI-PIK3CA), 170 (ARI-NABP2), and 49 (PIK3CA-NABP2), with 26 motifs shared across all three introns.

To experimentally test this preference, we inserted the ARI between nucleotides 576 and 577 of the GFP coding sequence and mutated the endogenous guanine at position 576 to adenine (A), thymine (T), or cytosine (C) (**Fig. 2B**). Quantitative PCR (qPCR) showed that all constructs produced equivalent levels of GFP mRNA; however, only the construct retaining G at position 576 yielded strong GFP protein fluorescence (**Fig. 3C**). Sanger sequencing of RT-PCR products confirmed that the 576A, 576T, and 576C mutants generated aberrant splice variants (**Fig. S2B-D**). In contrast, mutating the 3′ss region had no effect on splicing accuracy: all four nucleotides (A, T, C, and G) supported correct splicing (**Fig. 2D**). Together, these results establish a “G-ARI-N rule,” whereby a guanine at the −1 position relative to the 5′ss is essential for precise ARI splicing.

**Figure 3.**
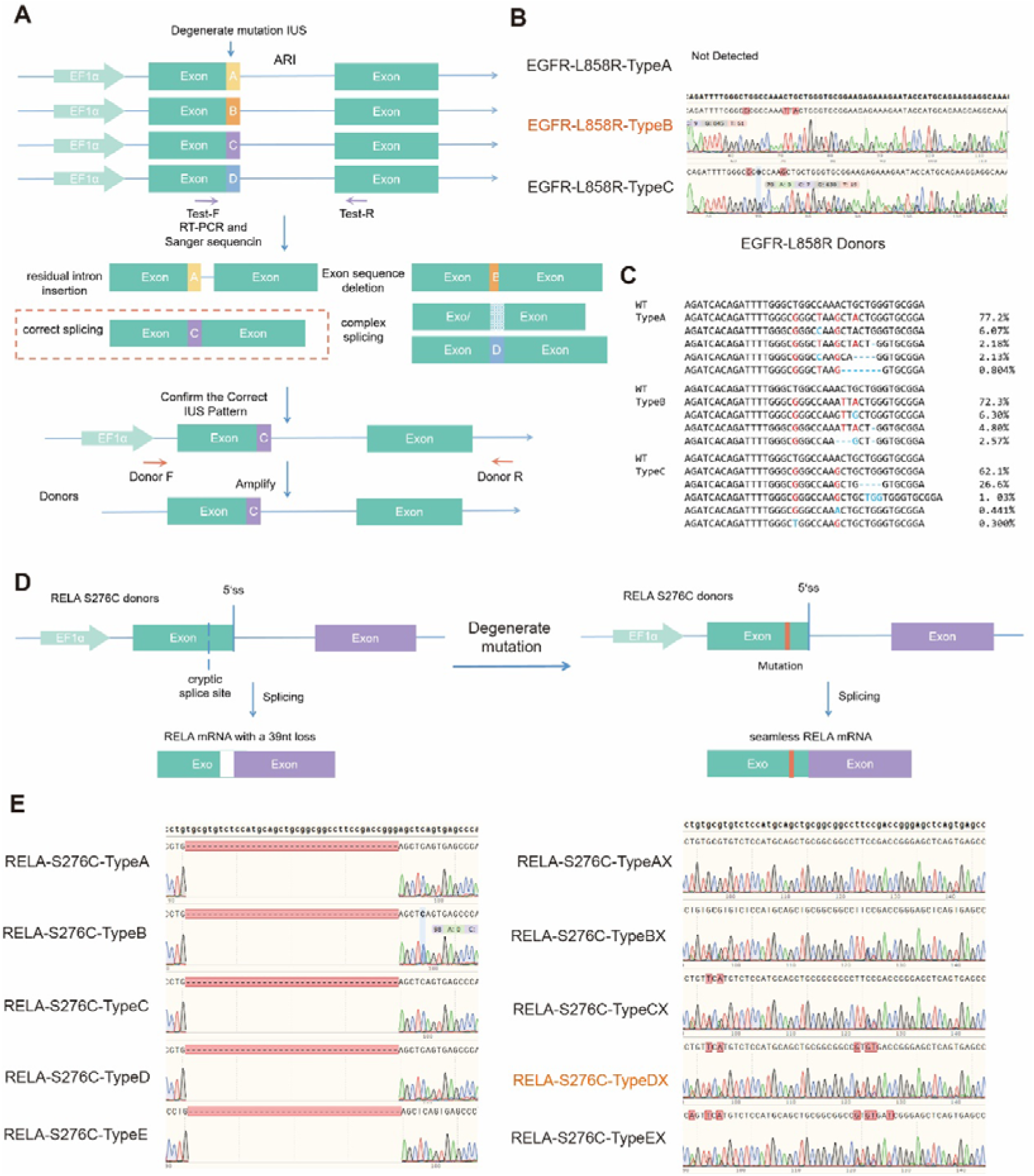
A. Workflow for selecting correct donors: (1) a series of donors with different degenerate IUS sequences were synthesized; (2) candidate donors were transiently transfected into target cells, followed by RT-PCR and Sanger sequencing; (3) the sequencing chromatogram was used to identify the correct donor. Only the correctly spliced donor in reporter assays are used for HDR. B. Chromatograms of RT-PCR products from EGFR-L858R candidate donors. C. NGS analysis of RT-PCR products from EGFR-L858R candidate donors. D. Illustration of a cryptic splice site identified in RELA donors, and degenerate mutations introduced to block its activation. The schematic compares RELA S276C donors driven by EF1α: the native construct utilizes a cryptic splice site near the 5’ss, resulting in RELA mRNA with a 39nt loss, whereas degenerate mutation at the site yields seamless RELA mRNA after splicing, preventing aberrant exon truncation. E. Chromatograms of RT-PCR products from RELA-S276C donors: the degenerate mutation at the cryptic splicing site restore the correct splicing in 5 tested donors.

To test the generality of this rule, we inserted the ARI into eight different locations within the GFP CDS, each adhering to the G-ARI-N rule. Of these, five constructs produced detectable GFP fluorescence, but only two (split-I and split-V) yielded correctly spliced mRNA transcripts (**Fig. S2F**). **The high frequency of mis-splicing likely reflects disruption of exonic splicing enhancers (ESEs)** (***14***)**, suggesting the need for locus-specific validation to ensure transcriptional stability.**

### The intron upstream sequences (IUS) influence RNA splicing

Given the influence of exonic splicing enhancers (ESEs), we hypothesized that local sequence context influences splicing accuracy. Because nucleotides downstream of the 3′ splice site (3′ss) showed no obvious preference (**Fig. 2A**), we reasoned that sequences upstream of the ARI 5′ss, termed intron upstream sequences (IUS), might contribute to aberrant splicing. To systematically evaluate the impact of IUS on ARI splicing, we designed a lentiviral vector library carrying a promoter-driven Flag-GFP fusion open reading frame (ORF) interrupted by a variable hexamer followed by a fixed guanine and the ARI (NNNNNNG-ARI). The GFP lacks a start codon; therefore, only when a given hexamer pattern permits correct ARI splicing can the reporter express Flag-GFP protein (**Fig. 2E**).

We screened this library to identify hexamer patterns that support efficient ARI splicing. HEK293T cells were transduced and selected with puromycin (0.7 μg/mL, 72 h) (**Fig. 2F**). Flow cytometry showed that 39.9 ± 0.3% of cells were GFP-positive (**Fig. 2G, S2G**). Genomic DNA from sorted GFP-positive and GFP-negative populations was extracted; the hexamer region was PCR-amplified and sequenced by NGS. Enrichment of each hexamer was calculated as its ratio in the GFP-positive versus GFP-negative library. Of 2,145 hexamers, 834 (≈40%) showed enrichment >2, indicating promotion of ARI splicing (**Fig. 2H, Table S1**).

We next asked whether these 834 hexamer patterns also promote splicing in the context of other introns. We replaced the ARI with introns derived from *NABP2*and *PIK3CA*and repeated the screen. The *NABP2*and *PIK3CA*libraries yielded 41.87 ± 0.45% and 50.1 ± 1.13% GFP-positive cells, respectively (**Fig. 2G, S2G**). The overlap among enriched hexamer patterns for the three introns is shown in a Venn diagram (**Fig. 2I**). These results suggest that although the influence of IUS on splicing may be a general phenomenon, the specific hexamer patterns that promote correct splicing differ substantially across introns.

Thus, there appears to be no universal IUS motif that guarantees accurate splicing, and ensuring proper ARI splicing cannot be achieved simply by substituting one artificial intron for another. Moreover, considering longer IUS regions (>6 nt) and diverse genomic insertion sites, the regulatory landscape is likely even more complex. **Unless splicing precision is rigorously validated, neither the robustness of this system nor the reliability of intron-based applications can be assured.**

### Identifying locus-specific IUS patterns using reporter vectors

To address aberrant splicing caused by ARI insertion, we systematically analyzed the influence of intron upstream sequences (IUS). Screening revealed that 38.9% (834/2,145) of hexamer patterns promoted correct ARI splicing, with varying rates for *NABP2* (52.2%, 905/1,738) and *PIK3CA* (11.1%, 297/2,673). Because ∼40% of hexamer patterns supported correct splicing, **we hypothesized that synonymous IUS mutations, which do not alter codons, might also permit accurate splicing.** To test this, we constructed a series of IUS-degenerate donor vectors under the EF1α promoter (**Fig. 3A**). RNA transcribed from these vectors recapitulated splicing, enabling RT-PCR and Sanger sequencing to identify correctly spliced donors.

For EGFR-L858R editing, three donor templates (Types A–C) were designed according to the G-ARI-N rule. Sanger sequencing confirmed that Type B generated the highest proportion of correctly spliced transcripts (**Fig. 3B**). To further characterize splicing outcomes, cDNA products were analyzed by high-throughput sequencing. Results showed that Types A and B produced >70% correct EGFR-L858R transcripts, with each mis-spliced product accounting for <10% of total reads, suggesting they likely arose from PCR artifacts. Type C, however, contained a minor variant comprising 26.6% of total reads, indicative of an aberrant splicing product. We therefore selected EGFR-L858R-TypeB as the donor for subsequent editing (**Fig. 3C**). Using this reporter assay, we also identified PIK3CA-H1047R-TypeC and FANCF-L15P-TypeA as functional donors (**Fig. S3 A, B**).

Surprising results emerged when we tested RELA-S276C donors: all candidates exhibited an identical 39-nt deletion in mRNA, suggesting activation of a cryptic splice site that removed 39 nt (*15, 16*). To restore correct splicing, we introduced degenerate mutations at the cryptic site. As expected, the modified RELA donors produced correctly spliced RNA, and we selected RELA-S276C-Dx as the donor.

Together, these results point to two major causes of aberrant splicing upon ARI insertion: (1) suboptimal IUS that perturbs splicing regulator recruitment and splice-site recognition, and (2) activation of cryptic splice sites leading to exon skipping. Both issues can be mitigated by introducing degenerate mutations into the donor sequence. Although splicing regulation is inherently complex, our approach that combining degenerate-mutated donor templates with RT-PCR and Sanger sequencing provides an efficient screening strategy to identify HDR donors that preserve correct splicing. This workflow represents a practical and robust pipeline to minimize splicing-related failure and may be broadly applicable to other engineered intron-based strategies (**Fig. S3C**).

### Establish ARI-HDR technology to generate gene edited cells

We applied the selected donor DNAs to edit RELA-S276C, EGFR-L858R, and PIK3CA-H1047R in HEK293T cells. Target-specific sgRNAs were designed, and HDR donors carrying the desired mutations were validated for correct splicing by RT-PCR and then amplified (**Fig. 4A**). Donor DNA and sgRNA-expressing lentiCRISPR-v2 vectors were transiently co-transfected into HEK293T cells, followed by puromycin selection (0.7 μg/mL, 72 h). DsRed-positive cells were isolated by FACS and expanded as single-cell clones. Clones were screened by PCR to confirm ARI integration and the presence of the targeted mutation. Successfully edited RELA-S276C, EGFR-L858R, and PIK3CA-H1047R cell clones were verified (**Fig. 4B**).

**Figure 4.**
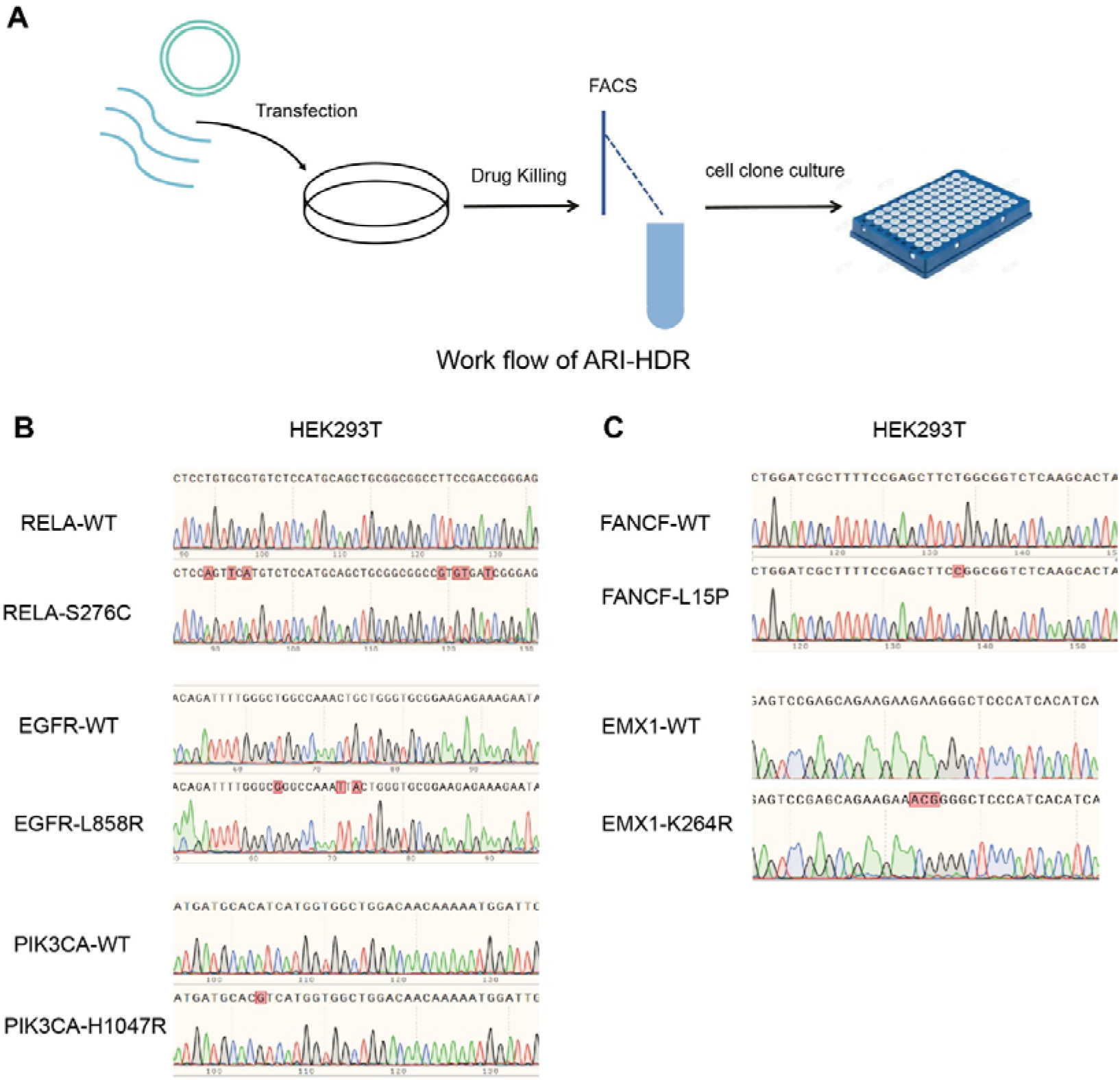
A. Workflow of ARI-HDR genome editing. The schematic illustrates the experimental process starting with plasmid transfection into cultured cells, followed by drug killing for selection, FACS (Fluorescence-Activated Cell Sorting) for screening, and subsequent cell clone culture in a multi-well plate to obtain edited clones. B. Sanger sequencing results of RT-PCR products from RELA-S276C–, EGFR-L858R–, and PIK3CA-H1047R–edited HEK293T cells. Chromatograms confirm the targeted mutations (RELA-S276C, EGFR-L858R, PIK3CA-H1047R) compared to their respective wild-type (WT) controls, with red boxes highlighting the specific nucleotide substitutions at the edited sites. C. Sanger sequencing results of RT-PCR products from FANCF-L15P– and EMX1-K264R–edited HEK293T cells. Sequencing traces verify the successful introduction of FANCF-L15P and EMX1-K264R mutations relative to WT, with red boxes indicating the altered nucleotides.

We also applied ARI-HDR to edit intron-less genes. Donors and sgRNAs were designed for EMX1-K264R and FANCF-L15P editing. Edited cells were successfully generated, and RT-PCR confirmed the presence of the expected mutated mRNA in all cases (**Fig. 4C**).

### Single-stranded DNA donors reduced the false-positive clones in HDR

Because the ARI contains a complete DsRed expression cassette, non-integrated or randomly integrated donors can produce DsRed signal, leading to false-positive cell clones (**Fig. 5A**). Both single-stranded DNA (ssDNA) and double-stranded DNA (dsDNA) donors can mediate HDR, but ssDNA typically achieves higher editing efficienc (*17–19*). Importantly, ssDNA cannot be transcribed in the absence of a promoter and is less susceptible to random integration via NHEJ (*20*).

**Figure 5.**
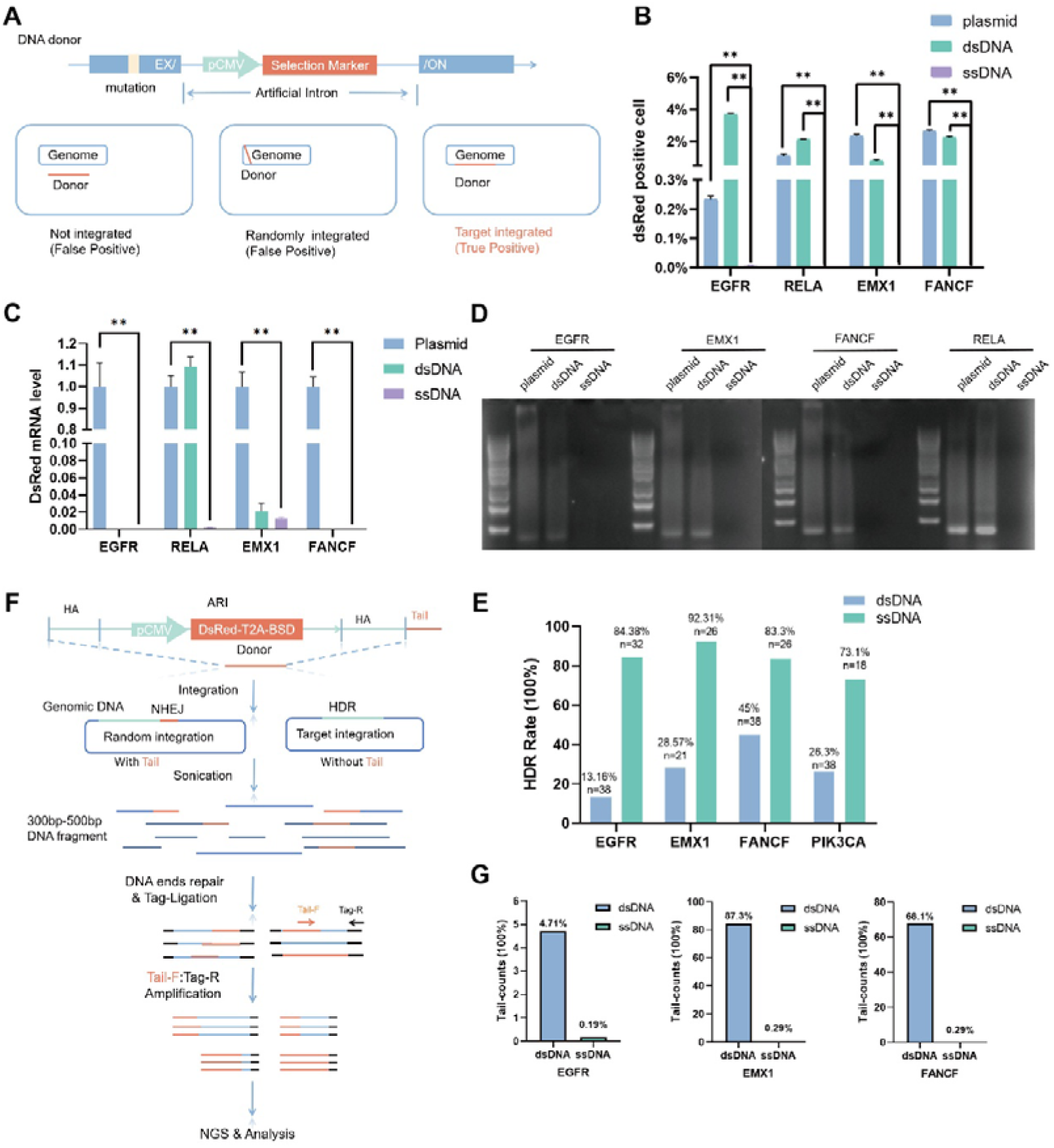
A. Graphic illustration of possible false-positive cells generated by ARI-HDR. Transient transfection or random integration of the donor may allow some cells to survive alongside truly edited cells, contributing to false-positive cells in FACS. B. Flow cytometry data from HEK293T cells transfected with plasmid, dsDNA, or ssDNA donors. The percentage of DsRed positive cells across four target loci (EGFR, RELA, EMX1, FANCF), comparing plasmid (blue), dsDNA (green), and ssDNA (purple) donor templates. Data are presented as mean ± SD from three independent experiments (n=3). Statistical significance was determined by unpaired two-tailed Student’s t-test (*p < 0.05, **p < 0.01). C, D. qPCR (C) and RT-PCR (D) analysis of DsRed mRNA from cells treated as in (B). The bar chart (C) shows the DsRed mRNA level across four target loci (EGFR, RELA, EMX1, FANCF), comparing plasmid (blue), dsDNA (green), and ssDNA (purple) donor templates. Data are presented as mean ± SD from three independent experiments (n=3). Statistical significance was determined by unpaired two-tailed Student’s t-test (*p < 0.05, **p < 0.01). Panel (D) displays the RT-PCR agarose gel electrophoresis results for the same four loci (*EGFR, EMX1, FANCF, RELA*), confirming the mRNA expression patterns observed in (C), with plasmid and dsDNA showing visible bands while ssDNA remains negligible. E. ARI-HDR editing efficiency mediated by dsDNA or ssDNA donors at the *EGFR, EMX1, FANCF*, and *PIK3CA* loci. The chart shows the HDR rate (100%) across these four target loci, comparing dsDNA (blue) and ssDNA (green) donor templates. Data are presented with sample sizes (n) indicated for each group (*EGFR*: dsDNA n=38, ssDNA n=32; *EMX1*: dsDNA n=21, ssDNA n=26; *FANCF*: dsDNA n=38, ssDNA n=26; *PIK3CA*: dsDNA n=38, ssDNA n=18). F. Workflow of TAIL-seq to detect the targeted and random integration of dsDNA or ssDNA donors: (1) genomic DNA from ARI-HDR edited cells mediated by ssDNA or dsDNA is extracted; (2) genomic DNA is sheared to 300-500 bp DNA fragments by sonication, followed by end-repair and tag-ligation; (3) TAIL-sequencing primers containing tag-sequences are used to amplify the DNA mixture (distinguishing HDR-mediated target integration without Tail and NHEJ-mediated random integration with Tail); (4) the amplicons are sent for NGS and analysis. G. Results of TAIL-seq in *EGFR, FANCF and EMX1* loci. Results were calculated as number of TAIL-containing reads normalized by primer-containing reads. Data are presented as percentage for dsDNA (blue) and ssDNA (green) groups. At the *EGFR* locus, dsDNA showed 37.91% (ssDNA: 1.88%); at *FANCF*, dsDNA showed 96.53% (ssDNA: 0.64%); at *EMX1*, dsDNA showed 99.22% (ssDNA: 0.61%).

To quantify the contribution of exogenous DNA format to false-positive signals, we transfected HEK293T cells with 0.5 μg of plasmid DNA, dsDNA, or ssDNA donors for RELA, EGFR, EMX1, and FANCF editing, without LentiCRISPR/gRNA plasmids. Flow cytometry at 36 h post-transfection detected DsRed-positive signals in plasmid DNA– and dsDNA-treated groups, whereas ssDNA donors showed no detectable DsRed (**Fig. 5B**). qPCR and RT-PCR for DsRed confirmed that no DsRed mRNA was transcribed from any of the DNA donors (**Fig. 5C-D, S4**). These results indicate that ssDNA does not generate false-positive edits.

To further evaluate the advantages of ssDNA as HDR donor, we performed ARI-HDR in *FANCF*, *EMX1*, *EGFR* and *RELA* loci using either ssDNA or dsDNA as donors. ssDNA up to 4000 nt in length were synthesized by Genscript. ssDNA donors yielded higher HDR efficiency than dsDNA donors across all tested loci (**Fig. 5E**). To assess random integration rates, we developed a TAIL-seq technique based on GUIDE-seq, in which only randomly integrated donors retain flanking TAIL sequences (*21*). Genomic DNA from ARI-HDR–edited cells (*EGFR*, *EMX1*, *FANCF*) were processed and sequenced (**Fig. 5F**). Compared to dsDNA, which harbored Tail-containing reads in all amplicons (*EGFR* 4.71%, *EMX1* 87.3%, and *FANCF* 68.1%), ssDNA donors yielded far fewer such reads (*EGFR* 0.19%, *EMX1* 0.29%, and FANCF 0.29%) (**Fig. 5G**). These results indicate that using single-stranded DNA as a donor leads to substantially lower random integration.

Taken together, these results demonstrate that ssDNA reduces false-positive events through two mechanisms: it cannot be transcribed, and it is less likely to be randomly integrated into the genome via NHEJ.

### ARI-HDR increases the editing efficiency by shortening the conversion tract

Another important question is what fraction of DsRed-positive cells carry the desired point mutation rather than merely an ARI knock-in (**Fig. 6A**). GEIS, a similar editing technique, have shown that conversion efficiency is inversely correlated with the distance between the target mutation and the DSB (the conversion tract) (*8*). To quantify this relationship, we designed donors with serial mutations (5–500 bp intervals) on homology arms for HDR at *RELA/EGFR/PIK3CA* loci. Genomic DNA from edited cell pools was PCR-amplified and subjected to Sanger sequencing. Statistical analysis revealed a significant negative correlation between conversion efficiency and distance, consistent with models predicting impaired resection kinetics at greater distances from the break (**Fig. 6B, C**) (*22*). These data indicate that minimizing donor-target distance enhances conversion efficiency.

**Figure 6.**
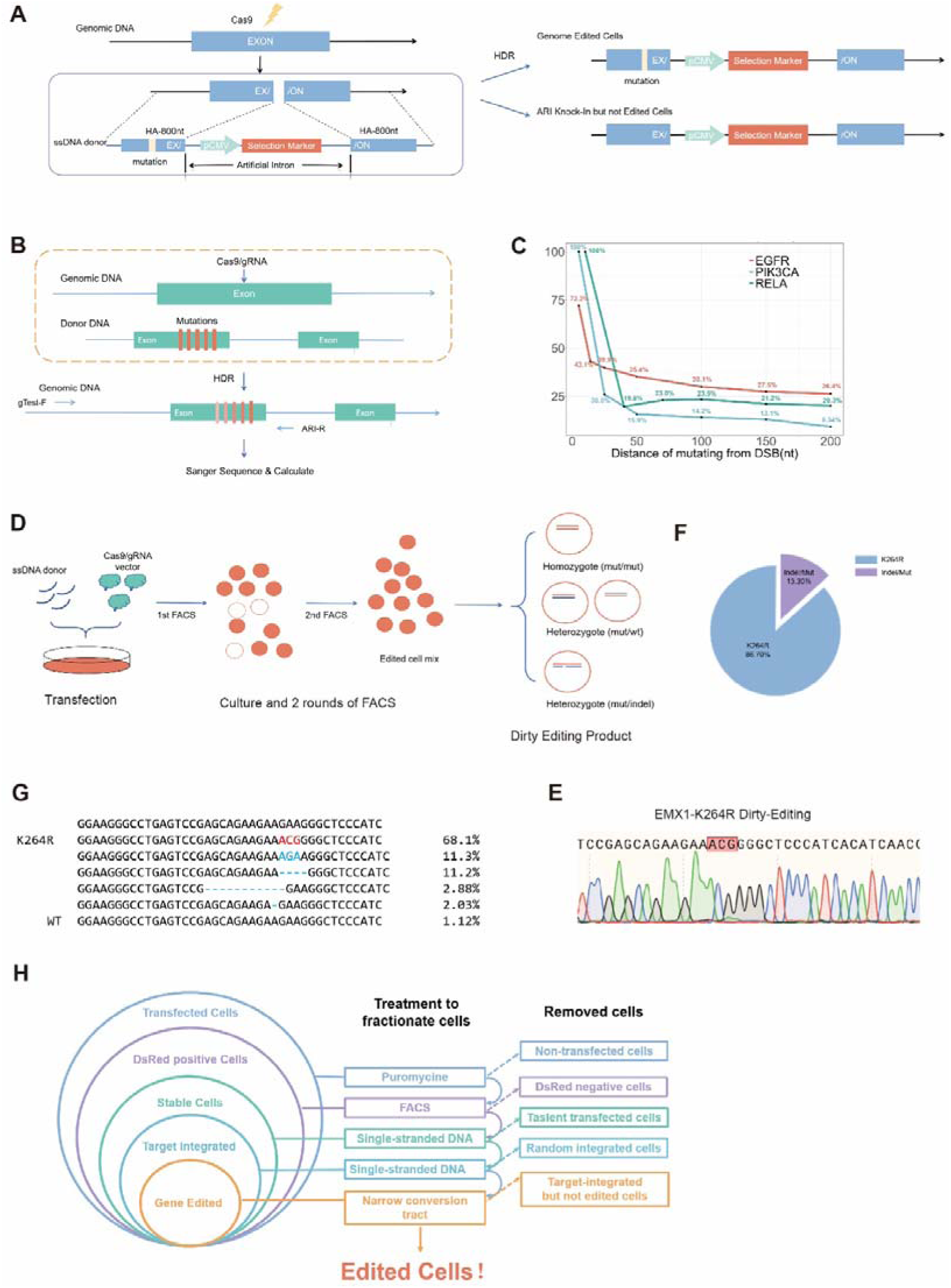
Proximity to the double-strand break (DSB) enhances gene conversion efficiency, and ARI-HDR strategy outperforms conventional GEIS by minimizing this distance. A. Illustration of ARI-HDR producing ARI Knock-in cells with or without target DNA editing. There is still a chance to produce false-positive cells in ARI-HDR, even after excluding random integration and transient expression of donors using ssDNA. B. Illustration of the experimental design for assessing the relationship between gene conversion efficiency and distance from the DSB. Donors with mutations at varying distances were co-transfected with Cas9/gRNA for HDR editing. PCR amplification (using gTest-F and ARI-R) followed by Sanger sequencing was performed. Conversion efficiency decreased as the distance between the DSB and the editing site increased across the three tested loci (*EGFR, PIK3CA, RELA*). C. Results from B. illustrate that gene conversion efficiency declines as the distance from the DSB increases across *EGFR*, *PIK3CA,* and *RELA*. ARI-HDR achieves higher efficiency by minimizing this distance. D. Workflow of dirty editing. Cells were co-transfected with ssDNA donor and Cas9/gRNA vector. Following transfection, cells underwent culture and two rounds of FACS, yielding an edited cell mixture. As predicted, the harvested mixture should consist of edited cells, including homozygote (mut/mut) and heterozygotes (mut/wt or mut/indel). E. cDNA analysis and G. NGS analysis of *EMX1-K264R* dirty editing products. Sanger sequencing (E) confirms the targeted *K264R* mutation (highlighted ACGG), while NGS quantification (G) shows the edited population comprises 73.04% K264R. F. Quantification of 500 amplicons from K264R-edited cells, and 86.7% are edited cells, 13.3% are indel/Mut. H. Summary of risks and corresponding mitigation strategies in ARI-HDR editing. The workflow illustrates a sequential cell fractionation approach starting from transfected cells. Treatment with puromycin selects for DsRed-positive stable cells, followed by FACS to exclude non-transfected, DsRed-negative, and transiently transfected cells. Subsequent selection using single-stranded DNA eliminates random integrated cells, ultimately isolating gene-edited cells via a narrow conversion tract while removing target-integrated but non-edited cells.

Unlike GEIS, which requires intron-targeting gRNAs, ARI-HDR allows gRNAs to target exons adjacent to the desired mutation, enabling a shorter conversion tract. To verify this advantage, we performed HDR using both strategies at the *EGFR* and *RELA* loci. ARI-HDR achieved higher conversion efficiencies than GEIS: 99.6% versus 69.7% for *RELA-S276C*, and 62.9% versus 45% for *EGFR-L858R*. (**Fig. S5A, B**). Thus, by enabling flexible sgRNA placement near the target site, ARI-HDR minimizes the conversion tract and improves editing efficiency.

### Dirty editing strategy: A Single-cell culture-free gene editing approach

After optimizing the ARI-HDR system to eliminate false-positive cells and enhance conversion efficiency, we obtained a mixed population of precisely edited cells containing various genotypes (homozygous, heterozygous, and monoallelic edits). We term this streamlined workflow dirty editing **Dirty Editing** (**Fig. 6D**).

To validate this approach, we applied dirty editing to introduce the *EMX1-K264R* mutation in HEK293T cells. After two rounds of FACS enrichment, the bulk cell population was harvested. RT-PCR amplification of *EMX1* cDNA followed by Sanger sequencing revealed that the vast majority of transcripts carried the *K264R* mutation (**Fig. 6E**). To quantify the editing efficiency, 500 individual amplicons were subjected to Sanger sequencing. Among these, 399 (79.8%) were EMX1-K264R mutants, 61 (12.2%) contained indels or other mutations, and 40 (8.0%) represented empty vectors, corresponding to 86.7% of EMX1 mRNA being correctly edited (**Fig. 6F**). Independent deep sequencing (NGS) of the same amplicons showed that 68.1% of total reads aligned with the *EMX1-K264R* sequence, confirming dominant expression of the edited allele.

## Discussion

Selection marker-based genome engineering has proven highly effective for gene fusion labeling and conditional gene regulation. However, placing a marker to report internal exon edits—small insertions, deletions, or point mutations remains challenging. We addressed this by using an artificial reporter intron to split the target exon. As it undergoes self-excision after splicing, keeping the corresponding mRNA scarless, the ARI can be inserted into virtually any exon, solving the marker placement problem. Given that splicing machinery is conserved across eukaryotes, this strategy holds broad potential for genomic manipulation (*12*).

Splitting an exon is inherently risky. Previous attempts at artificial intron–based gene editing have been reported (*23, 24*) but proved difficult to replicate at other loci (*10*), often causing indels in the transcript and unintended gene knockout. This study identified two principal causes: the influence of intron upstream sequences (IUS) on splicing accuracy, and the activation of cryptic 5′ splice sites upon intron insertion. Based on our findings, the degenerate-mutated donors combined with assaying the splicing products could address both problems. **This approach might resolve the vast majority of mis-splicing events.** More broadly, the identification of IUS as a universal splicing determinant reveals a previously overlooked mechanism of gene disruption: **exonic mutations can trigger functional knockout through splicing dysregulation, a pitfall that conventional ectopic expression assays are ill-suited to detect.** This mechanism may explain a subset of disease-associated synonymous or missense mutations whose pathogenicity has remained mechanistically unclear(*25*).

Transient expression and random integration of reporter genes via NHEJ or LIG4-independent mechanisms such as SINE-mediated integration remain major sources of false-positive cells in selection-based editing (*26*). Recent advances highlight single-stranded DNA (ssDNA) as an optimal donor for HDR, owing to its structural simplicity, low immunogenicity, and enhanced recruitment by RAD51 through endogenous repair pathways (*18, 27*). Enzymatic DNA synthesis now enables scalable production of long ssDNA (up to 7 kb) with >98% accuracy (*28*). In this study, we used 4-kb ssDNA as the HDR donor. Our data show that ssDNA cannot be transcribed but can mediate HDR, and TAIL-seq confirms that its random integration is dramatically reduced. Consequently, ssDNA effectively excludes false-positive cells, yielding targeted HDR rates of up to 90%.

A gene conversion tract refers to the segment of DNA transferred from a donor to an acceptor sequence during homologous recombination (*29*). Conversion efficiency depends critically on the proximity of the DSB to the editing site: it remains 70–99% when the distance is ≤10 nt but declines sharply beyond 50 nt (*8, 22*). The ARI-HDR system optimizes this by strategically positioning intronic sequences to minimize the DSB-to-edit distance. Indeed, when the conversion tract was shorter than 5 nt, efficiency reached nearly 100% for both RELA and PIK3CA.

Building on the high HDR efficiency achieved with ssDNA-mediated ARI-HDR and the enhanced conversion efficiency from shortened conversion tracts, we developed a “Dirty-Editing” workflow that bypasses laborious single-cell culture and genotyping (*30*). For EMX1-K264R editing in HEK293T cells, where the ARI is only 5 nt from the DSB, edited mRNA accounted for over 85% of EMX1 transcripts. This workflow is particularly suited for cell types in which single-cell cloning is impractical.

Several limitations remain. Non-coding genes cannot be analyzed by degenerate mutation assaying, making it difficult to apply ARI-HDR for accurate editing in these contexts. The current workflow relies on FACS, limiting applicability to shear-sensitive cell types. Long ssDNA synthesis remains costly, though advancing enzymatic methods may reduce this barrier. Performance in hard-to-transfect cells such as primary T cells or iPSCs requires further validation. Looking forward, multiplexed ARI cassettes with distinct fluorescent markers could enable multi-locus editing. Coupling with viral or lipid nanoparticle delivery may extend ARI-HDR to in vivo settings. The ARI module could also be integrated into base or prime editors to broaden enrichment beyond classical HDR. Addressing these limitations will be key to translating this platform into broader applications.

Among genome editing approaches, homology-directed repair (HDR) offers unmatched precision for targeted insertions, deletions, or substitutions with minimal off-target effects (*31*). To overcome its low efficiency, selection marker–based enrichment has been widely used to isolate edited clones. Here, we present an efficient HDR strategy that uses an artificial reporter intron to label edits. Because splicing is conserved across eukaryotes, this strategy is in principle applicable to any coding gene in any eukaryotic system. Under optimized conditions, editing efficiency exceeded 86%.

This performance was achieved through a combination of complementary measures: puromycin selection to eliminate untransfected cells, FACS enrichment based on the ARI-encoded fluorescent marker, the use of ssDNA donors to suppress false positives from transcription and random integration, and strategic placement of the Cas9 cut site adjacent to the ARI to favor true editing over marker-only integration (**Fig. 6H**). Together, these measures transform HDR from a low-efficiency, locus-restricted tool into a robust and generalizable platform for precise genome engineering.

## AUTHOR CONTRIBUTION

S.W. conceived the study, performed most of the experiments, and wrote the manuscript. W.H. provided the majority of the funding. X.W. and W.H. provided the research platform. M.S. participated in some of the experiments. K.W. performed the statistical analysis. M.Z. edited the manuscript.

## Acknowledgement

We thank Mr. Wugen Zhan for His professional aid in FACS experiment. We thank SHENZHEN SYNTHETIC BIOLOGY INFRASTRUCTURE and Core Facility of Shenzhen Institute of Synthetic Biology offering platform for experimental support. We thank Dr. Yuanzhong Wu from Sun Yat-sen University Cancer Center for his constructive advice on experimental design and manuscript preparation.

---

**Figure S1.**
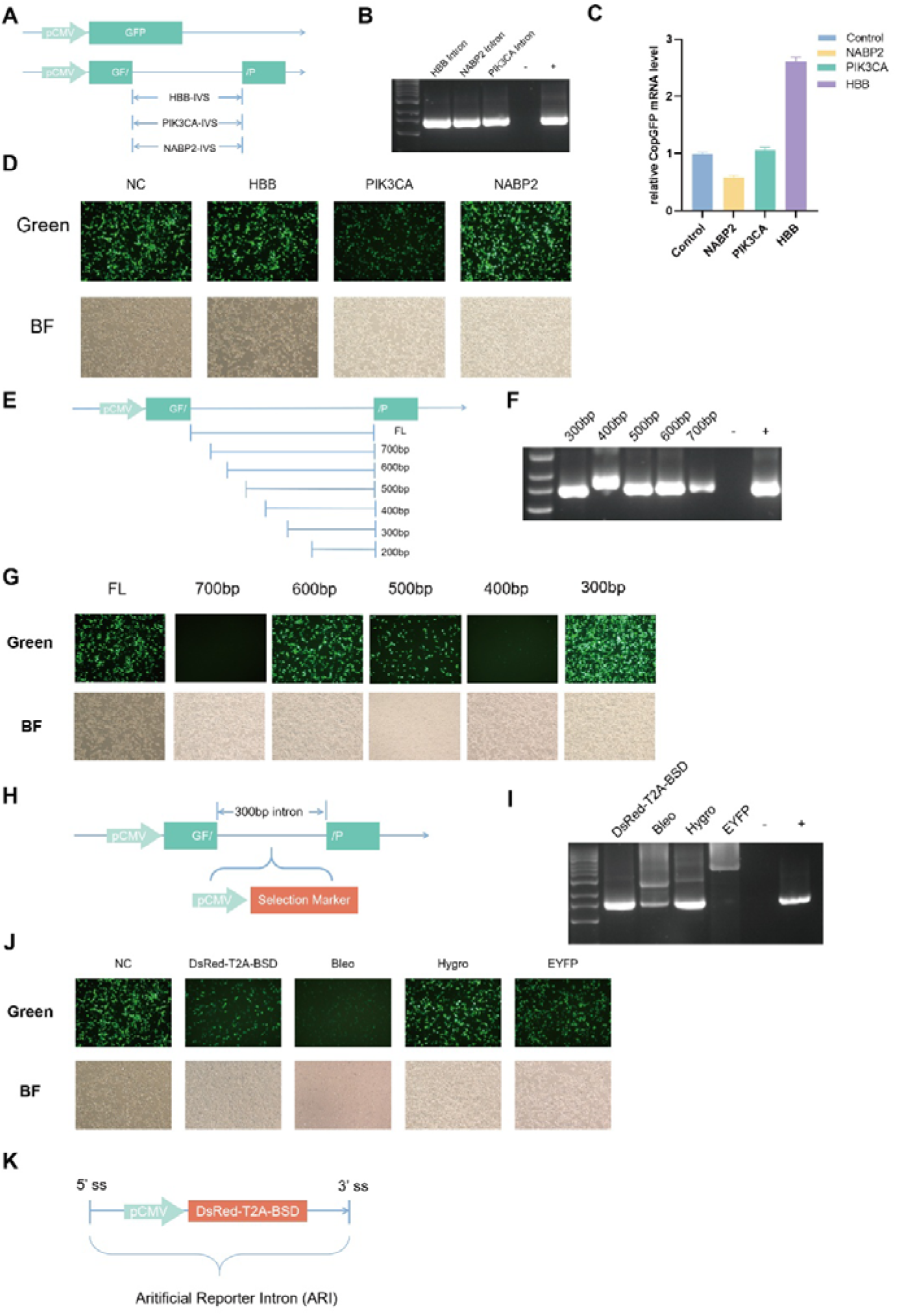
Process to select and optimize the artificial reporter intron (ARI). **A.** Split-GFP constructs (pCMV-GF-/P) carrying introns from *HBB*, *PIK3CA*, or *NABP2*. **B-D.** All introns spliced correctly by RT-PCR (B); *HBB* intron yielded the highest CopGFP mRNA by qPCR (C); GFP reconstitution was confirmed by fluorescence imaging (D). **E-G.** Truncated *HBB* introns (full-length to 300 bp) were tested (E); only 300-, 500-, and 600-bp variants supported correct splicing by PCR (F) and fluorescence (G). **H-J.** Different selection markers were inserted into the 300-bp intron (H); only the DsRed-T2A-BSD cassette avoided undesired splicing (I) and enabled correct splicing with detectable fluorescence (J). **K.** Final ARI design: a 300-bp *HBB*-IVS containing DsRed-T2A-BSD, flanked by 5′ and 3′ splice sites.

**Figure S2.**
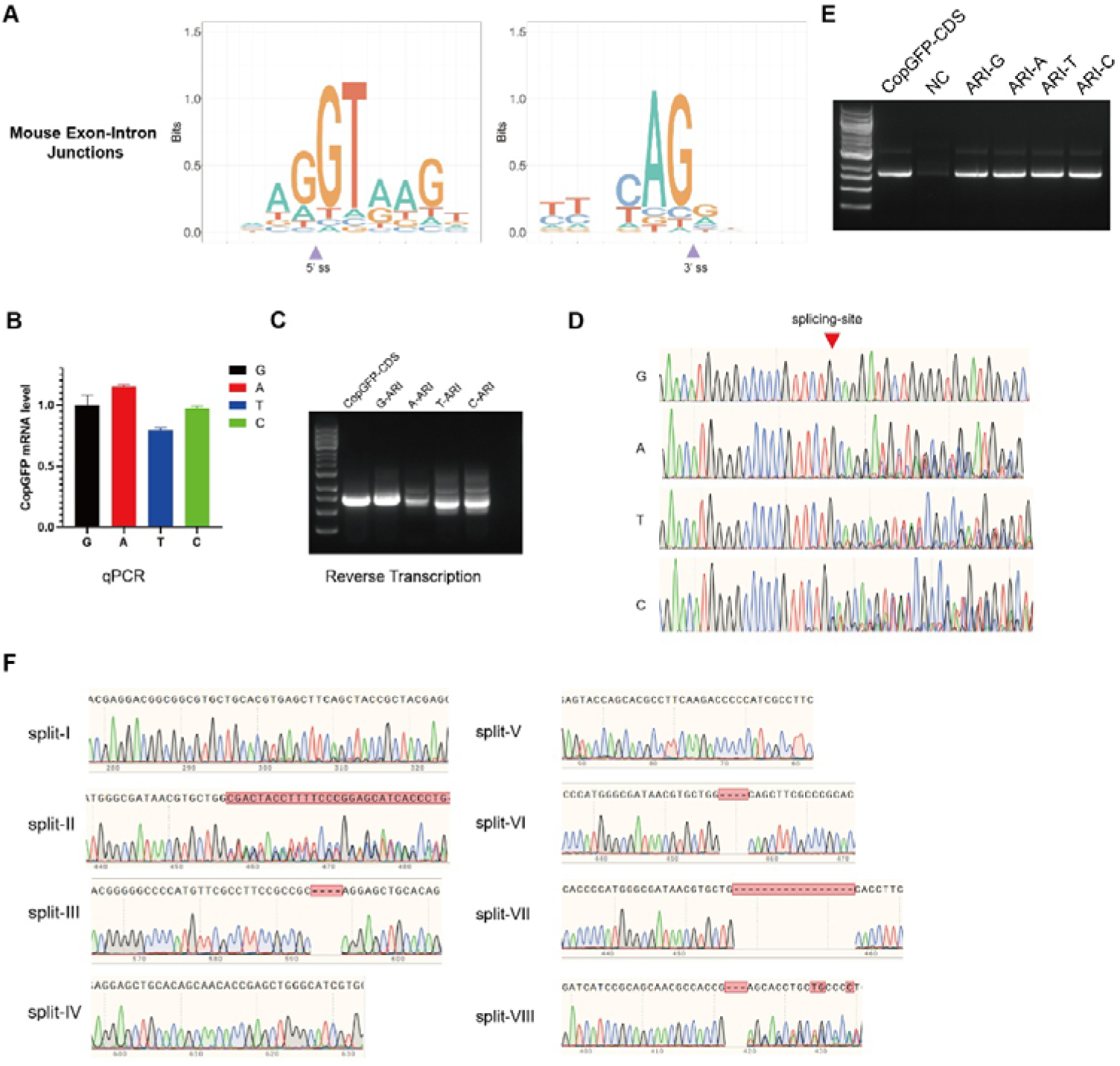
A. Sequence logo analysis of mouse exon-intron junctions at 5′ and 3′ splice sites. **B-D.** Expression of a CMV-driven GFP ORF disrupted by ARI insertion with degenerate mutations in the 5′ flanking region was assessed by qPCR (B), RT-PCR (C), and Sanger sequencing confirming the splicing site (D). **E.** RT-PCR analysis of the reporter GFP CDS interrupted by ARI with degenerate mutations in the 3′ flanking regions (CopGFP-CDS, NC, ARI-G, ARI-A, ARI-T, ARI-C). **F.** Sanger sequencing chromatograms of GFP CDS split by ARI at eight different positions (split-I to VIII) following the G-ARI-N rule, showing various splicing outcomes.

**Figure S3.**
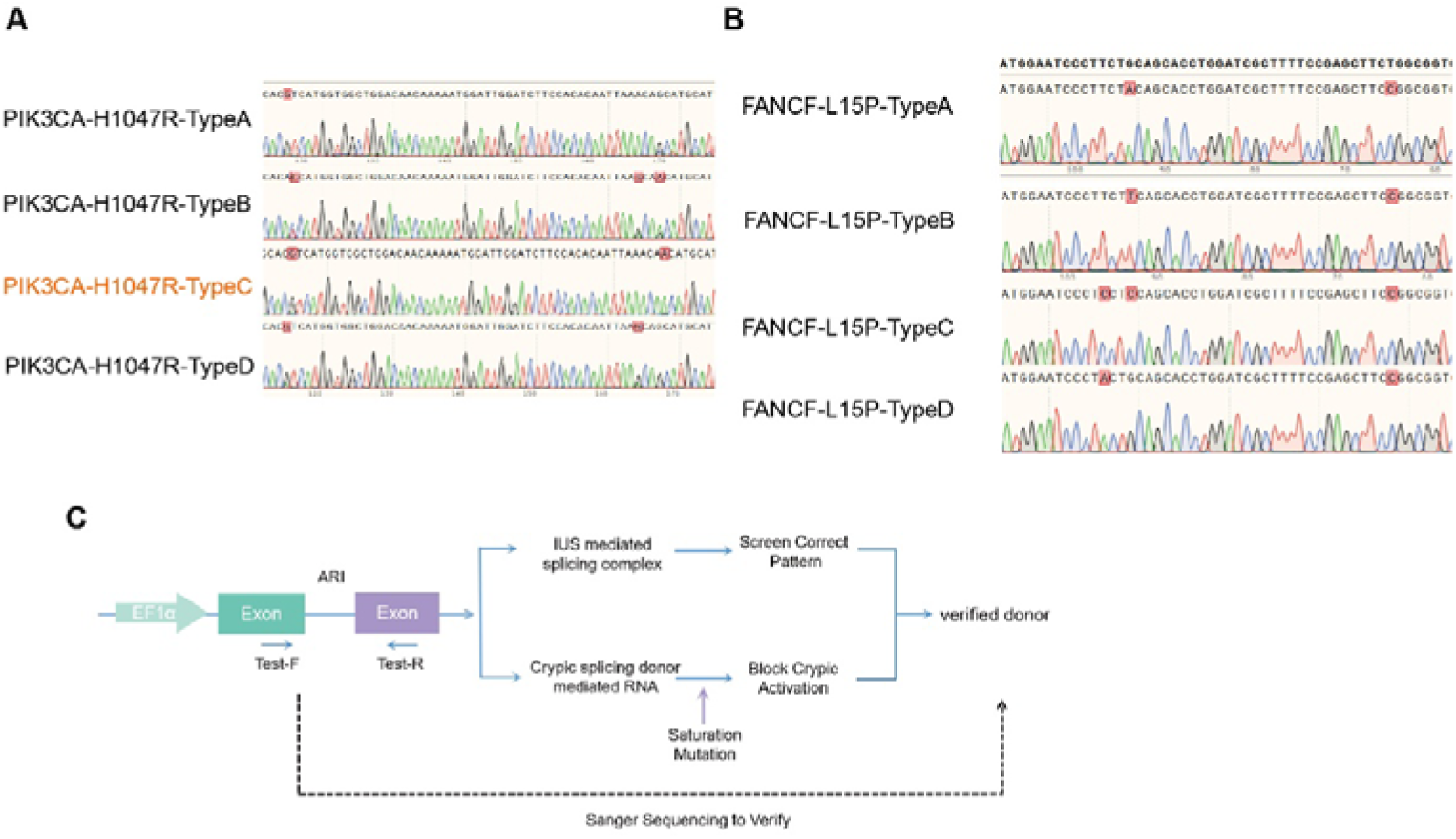
A. Sanger sequencing of RT-PCR products from *PIK3CA-H1047R* donors with IUS degenerate mutants. Types B, C, and D showed no aberrant splicing, and Type C was selected as the donor for ARI-HDR. **B.** *FANCF-L15P* donors assay showed all variants (Type A–D) produced no aberrant splicing. **C.** Schematic of the donor screening workflow. An EF1α-driven exon-ARI-exon construct is evaluated via Test-F/R primers. Transfection and RT-PCR/Sanger sequencing reveal splicing products. Undesired variants from IUS are identified via degenerate mutants, and cryptic splicing activation is blocked by site-specific degenerate mutation to verify the donor.

**Figure S4.**
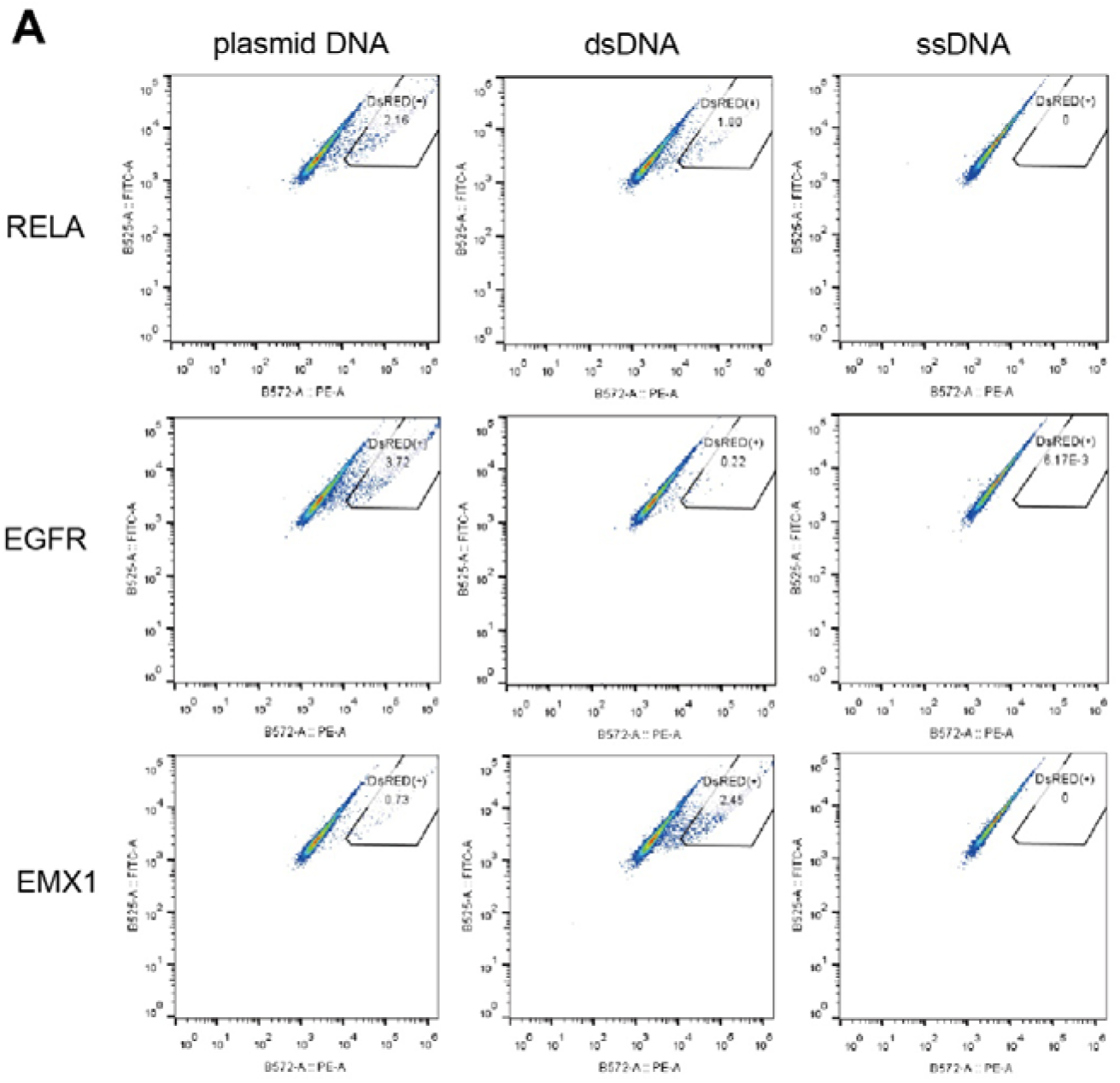
Flow cytometry analysis of HEK293T cells transfected with plasmid DNA, dsDNA, or ssDNA.

**Figure S5.**
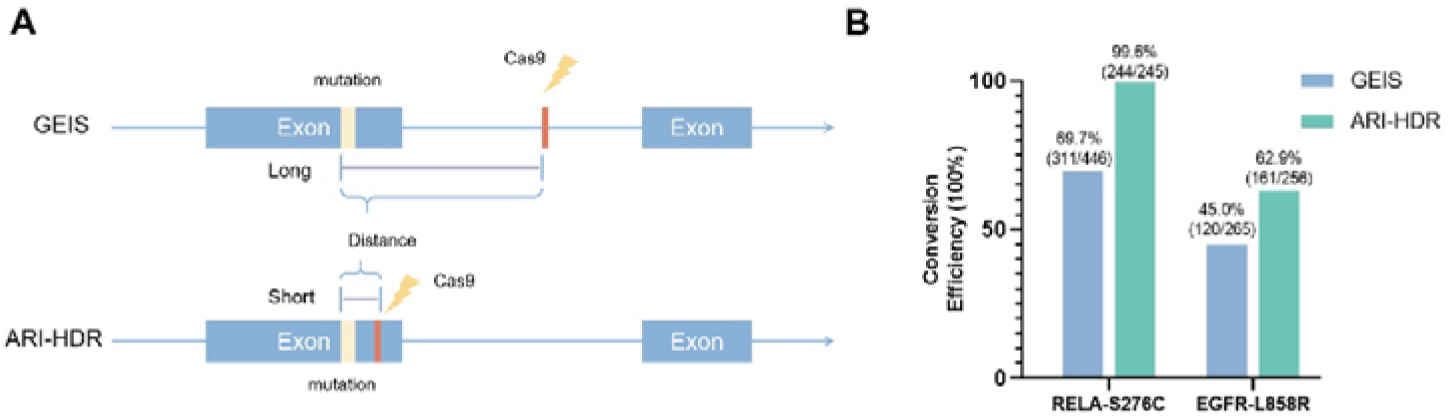
Comparison of GEIS and ARI-HDR strategies for locus editing. **A.** Schematic illustrating the mechanistic differences between GEIS and ARI-HDR. ARI-HDR achieves higher conversion efficiency by shortening the conversion tract. **B.** Comparison of editing efficiency at the same loci. For *RELA-S276C*, ARI-HDR achieved 99.6% (244/245) versus GEIS 84.0% (194/231); for *EGFR-L858R*, ARI-HDR achieved 78.6% (162/206) versus GEIS 55.0% (126/229). ARI-HDR consistently outperformed GEIS across both targets.

## Methods and materials

### Plasmids and donors

DSBs were generated by CRISPR–Cas9 technology with LentiCRISPR-V2 (Addgene, #98290) carrying the indicated intron-targeting sgRNAs (RELA: 5’-UCA CUG AGC UCC CGG UCG GA-3’, EGFR: 5’-GCU GGC CAA ACU GCU GGG UG-3’, PIK3CA: 5’-AUG CAU GCU GUU UAA UUG UG-3’, EMX1: 5’-GAG UCC GAG CAG AAG AAG AA-3’, FANCF: 5’-GGA AUC CCU UCU GCA GCA CC-3’).

The pUC19-EF1-ARI construct were used for donor synthesis. Primers (Donor-F: 5’-ACC GGC GCC TAC GGT ACC GGA T-3’, Donor-R: 5’-CCG GAG ACG GTC ACA GCT TG-3’) were used to amplify the donors. Detailed donor sequences for editing were in supplementary materials.

Single-stranded DNA were custom made by GenScript (China).

### Cell culture and FACS

HEK293T cells were obtained from the American Type Culture Collection (ATCC; CRL-3216). Cells were cultured in DMEM (Dulbecco’s modifed Eagle’s medium) containing 10% fetal bovine serum with 1% penicillin-streptomycin at 37 °C and 5% CO2.

HCT116 cells were obtained from Ubigene (China), cultured in RPMI 1640 (Roswell Park Memorial Institute 1640) containing 10% fetal bovine serum supplemented with 1% penicillin-streptomycin at 37 °C and 5% CO2.

Fifteen micrograms of LentiCRISPR-V2 and 2 µg of donor DNA were co-transfected into 5 × 10^6^ cells with Lipofectamine 3000 (Thermo Fisher Scientific) according to the manufacturer’s instruction. Transfected cells were treated with 0.6 µg/mL puromycin for 72 h followed by 48h culture with complete media. Cells were digested with 0.5% of Trypsin, resuspended in PBS and filtered in 40 µm cell strainer (BD Falcon). The 1^st^ round of FACS were performed in BD AriaIII to enrich the dsRed(+) cells, and followed by a complete media culture supplemented with Y27632 (Abclone) for 2 weeks. The 2^nd^ round of FACS were processed to select single-cell clone for single-cell spanning in 96-well plates.

### Genomic DNA extraction and analysis

Genomic DNA was extracted using the MolPure® Cell/Tissue DNA Kit (YEASEN,18700ES70). PCR of single clones derived from genomic DNA was processed to verify that the clone possessed the desired sequence. PCR was performed for 35 cycles on an Eppendorf thermocycler with denaturation at 94 °C for 15 s, annealing at 58 °C for 15 s, and extension at 72 °C for 90 s using 2× Hieff Canace® Gold PCR Master Mix (YEASEN, 10149ES01).

Primer pairs used for cloning validation are:

RELA:(RELA-gTEST-F: GGC ACC CAC TGG ACT A, pUni-R: GTA AGT TAT GTA ACG CGG AA),
EGFR: (EGFR-gTEST-F: ATA AGA AGT CTG CAG AAC TT, pUni-R: GTA AGT TAT GTA ACG CGG AA),
PIK3CA: (PIK3CA-gTEST-F: GAG GAA AGT CAG TCA ACC AT, pUni-R: GTA AGT TAT GTA ACG CGG AA),
EMX1: (pUni-F: CGG CAG TTG GGA TTC GTG AA, EMX1-gTEST-R: CTC CCT CTC CCA TTG AAC TA),
FANCF: (pUni-F: CGG CAG TTG GGA TTC GTG AA, FANCF-gTEST-R: ACT TTA GTG ACT AGC CGC CAC).

### Conversion efficiency detection

Multiple DNA variations on the left HA for RELA, PIK3CA and EGFR GEIS were introduced to HEK293T cells by ARI-HDR. FACS were processed to enrich ARI-HDR edited cells. Genomic DNA was subsequently extracted from the sorted population. The integrated DNA fragment within the homology arms was amplified by PCR. The resulting amplicons were then cloned into an EcoRI-linearized pCDH-cmv-mcs-EGFP-puro vector for transfection. Finally, individual bacterial colonies were subjected to Sanger sequencing. The editing efficiency was calculated as the ratio of each specific mutation within the total sequenced clones.Primers used for clone were:

RELA: (arm-RELA-gTEST-F: <u>AAG ATT CTA GAG CTA GCG</u> GGC ACC CAC TGG ACT A, arm-pUni-R: <u>GGATCCGATTTAAATTCG</u> GTA AGT TAT GTA ACG CGG AA),
EGFR: (EGFR-gTEST-F: <u>AAG ATT CTA GAG CTA GCG</u> ATA AGA AGT CTG CAG AAC TT, pUni-R),
PIK3CA: (PIK3CA-gTEST-F: <u>AAG ATT CTA GAG CTA GCG</u> GAG GAA AGT CAG TCA ACC AT, pUni-R),

#### TAIL-Seq

This experiment was designed based on guide-seq (*21, 32*).

ARI-HDR processed cells were harvested for genomic DNA extraction. 100μg of Genomic DNA were subjected to sonication to generate 300-500bp small fragments. Fragmented DNA were purified, end-repaired and tag-ligated.

Primer TAIL-F1: 5’-TGA CGC GCC CTG ACG GGC TT-3’, and Tag-R: 5’-TCT TCC

TAA GAC CGC TTG G-3’ were used to PCR amplify in thermocycler.

Three 20 μL PCR reactions were set up as follows:

10 μL 2× PCR amplification buffer
0.5 μL TAIL-F1 (10 μM, final concentration 0.25 μM)
0.5 μL Tag-R (10 μM, final concentration 0.25 μM)
333 ng template fragmented DNA.
Nuclease-free ddH□O added to 20 μL.

Thermal cycling conditions: 94°C 3 min pre-denaturation; 30 cycles of 94°C 15 s, 55°C 15 s, 72°C 30 s; final 72°C 5 min extension, hold at 4°C.

PCR products were purified and followed by an additional PCR reaction using primers TAIL-F2: 5’-CAA CTC CTT GGC TCA CAG AAC GAC ATG GCT ACG ATC CGA CTTTCT GCT CCC GGC ATC CGC TT-3’, and Tag-R. Thermal cycling conditions: 94°C 3 min pre-denaturation; 20 cycles of 94°C 15 s, 55°C 15 s, 72°C 30 s; final 72°C 5 min extension, hold at 4°C. PCR product were sent BGI for NGS.

### Intron upstream sequence library detection

We constructed lentiviral reporter vectors (psin-reporter-Flag-MCS-GFP) in which the coding sequence (CDS) of FLAG tag was fused in-frame to the CDS of GFP via a multiple cloning site (MCS) linker. Notably, the translation of GFP relies on the start codon (ATG) provided by the FLAG tag sequence. Three distinct libraries were generated by inserting sequences (6N-G-ARI, PIK3CA-IVS, EGFR-IVS) into the MCS. Each library was individually packaged into lentiviral particles. Subsequently, HEK293T cells were infected with the respective viruses at a low multiplicity of infection (MOI < 0.3). To enrich successfully transduced cells, culture medium was supplemented with 0.6 μg/ml puromycin at 48 hours post-infection. Following selection, GFP-positive and GFP-negative cell populations were isolated by fluorescence-activated cell sorting (FACS). Genomic DNA was then extracted from these sorted populations, and the integrated 6N region was amplified by PCR for subsequent next-generation sequencing (NGS) analysis.

### Bioinformatic analysis

All data processing and analysis were performed using Python v3.12 and R v4.5.2.

### Exon–intron junction sequence analysis

Exon–intron junction logos were built using the GRCh38 human reference genome and GENCODE annotation (v44). All exon records were parsed from the GTF file; for each exon, we defined an 11-bp window spanning the splice junction—4 bp upstream to 7 bp downstream (−4 to +7) for the 5′ss, and 7 bp upstream to 4 bp downstream (−7 to +4) for the 3′ss. BED intervals for the two classes of splice sites were generated separately. Genomic sequences were extracted with seqtk (v1.5) and merged with the exon annotation table. When an exon fell on the minus strand, the retrieved sequence was reverse-complemented to maintain the 5′→3′ orientation relative to the transcript. For each splice-site type, base frequencies (A, C, G, T) were tallied position by position across all oriented 11-mers and normalized by the total number of sites. These frequency matrices were then visualized as sequence logos using the R package ggseqlogo (v0.2.2), with the height of each nucleotide letter at each position scaled proportionally to its frequency.

### Data processing for screening 6N sequences

Illumina paired-end sequencing reads from the GFP-positive and GFP-negative cell populations were first demultiplexed and quality-filtered using Trimmomatic (v0.39) with default parameters for adapter removal and low-quality base trimming (leading/trailing quality < 20, sliding window 4:15). The hexanucleotide (NNNNNN) sequences immediately upstream of the G-ARI were extracted from the aligned reads. For each of the three independent intron contexts (TRI, NABP2, and PI3KCA), the raw count of each unique 6N motif was tallied separately for the GFP-positive (High) and GFP-negative (Low) libraries. To ensure statistical robustness, motifs with fewer than 10 counts in either the High or Low group were excluded from further analysis.

For each gene, the total number of reads mapped to all 6N motifs was computed for both groups (High_total and Low_total). The enrichment score for a given motif (i) was then defined as the log□-transformed ratio of its normalized counts in the two populations:

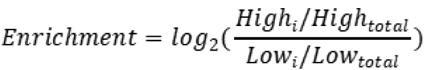

Motifs with higher enrichment values are preferentially associated with productive ARI splicing and GFP expression. After calculating enrichment scores for all qualifying motifs within each gene, the motifs were sorted in ascending order of their enrichment scores, and the resulting rank-ordered list was plotted as a scatter plot.

### Tail-seq data analysis

To evaluate the proportion of reads containing the Tail sequence motif in the Tail-seq NGS dataset, assembled reads were screened for the presence of a tail motif. The number of reads containing Tail-Sequence: ‘CAAGCTGTGAC’ was then normalized against the total number of primer containing reads.

### Reporter Assay to confirm splicing-guaranteed donor

To identify suitable donors that facilitate the desired splicing, a series of IUS degenerate mutant donors were synthesized. Donor plasmids were transfected into HEK293T cells. RNA extraction and RT-PCR were performed after 24 hours, and the products were subjected to paired-end NGS sequencing. The NGS data were analyzed as follows:

Step 1: The data were decompressed.
Step 2: The paired-end reads were assembled.
Step 3: Reads containing the X gene were filtered.
Step 4: Unique sequences longer than 50 nt were extracted from the FASTA file, and the copy number of each unique sequence was calculated and exported as a CSV file.

After removing endogenous variant, variants were listed according to their counts.

### Data Availability

The analysis scripts in this study can be found in a github repository: https://github.com/MedDataX/Precise-and-Effecient-Gene-Editing-via-Artificial-Reporter-Intron-Assisted-HDR.git

